# Non-contact microfluidic analysis of the stiffness of single large extracellular vesicles from IDH1-mutated glioblastoma cells

**DOI:** 10.1101/2022.08.19.504310

**Authors:** Mi Ho Jeong, Hyungsoon Im, Joanna B. Dahl

## Abstract

In preparation for leveraging extracellular vesicles (EVs) for disease diagnostics and therapeutics, fundamental research is being done to understand EV biological, chemical, and physical properties. Most published studies investigate nanoscale EVs and focus on EV biochemical content. There is much less understanding of large microscale EV characteristics and EV mechanical properties. We recently introduced a non-contact microfluidic technique that measures the stiffness of large EVs (>1 μm diameter). This study probes the sensitivity of the microfluidic technique to distinguish between EV populations by comparing stiffness distributions of large EVs derived from glioblastoma cell lines. EVs derived from cells expressing the IDH1 mutation, a common glioblastoma mutation known to disrupt lipid metabolism, were significantly stiffer than those expressed from wild-type cells. A supporting lipidomics analysis shows that the IDH1 mutation increases the amount of saturated lipids in EVs. Taken together, these data suggest that high-throughput microfluidics is capable of distinguishing between large EV populations that differ in biomolecular composition and therefore structure. These findings advance the understanding of EV biomechanics, in particular for the less studied microscale EVs, and demonstrate microfluidics to be a promising technique to perform clinical EV mechanophenotyping.

## 1. Introduction

Extracellular vesicles (EVs) are cell-derived, membrane-bound vesicles in heterogeneous subpopulations differentiated based on their biogenesis, sizes, density, and biomarkers. These include nanoscale vesicles (e.g., exosomes, 50 – 200 nm in diameter), microvesicles (200 – 1000 nm), and atypically large vesicles (e.g., large oncosomes, apoptotic bodies, giant extracellular vesicles, 1 – 5 μm). EVs carry various biomolecules, such as proteins, nucleic acids, and lipids, from their cellular origins and thus play important roles in cell-to-cell communications as molecular messengers by transferring these molecules to recipient cells. EVs are also attractive circulating biomarkers for molecular diagnosis of various diseases and carriers for therapeutics. Most current EV research focuses on molecular profiling of EVs and their biological roles and less on the physical characteristics beyond their sizes and concentrations. Better understanding of EVs’ physical properties will contribute to their characterization and use for diagnostic and therapeutic purposes.

Stiffness is a high-level indicator of biomolecular content and molecular arrangements (conformations). The stiffness of vesicles or cells sheds light on ongoing processes and is the end-result of many factors. Several studies in the cellular biomechanics literature have demonstrated that different expression levels of genes lead to different biomolecular content and ultimately different cellular behaviors. In one such example, researchers found that pancreatic cancer cell lines had different levels of genes, resulting in different structural protein contents and stiffnesses that impacted cellular invasion potential.^[1]^ The surrounding microenvironment can affect biomembrane and cell stiffness,^[2–5]^ and conversely, cells can modify the stiffness of their microenvironment.^[6]^ Even passive systems can experience mechanical changes due to the surrounding environment. For instance, the bending stiffness of synthetic giant vesicles can decrease by a factor of 4 as the concentration of sucrose in the surrounding aqueous solution is increased from 0 to 3 moles per liter.^[7]^ Cell and biomembrane mechanical stiffness can influence or be influenced by cellular processes.^[8]^ Therefore, stiffness has the potential to be a valuable metric to assess the range of the physical manifestations of cellular genetics, environmental conditions, and various cellular processes. Within the field of EVs, stiffness could add additional information to the conventional omics information (genome, transcriptome, proteome, lipidome) that are being used to distinguish EV subpopulations.

Recent atomic force microscopy (AFM) studies of EVs demonstrated the ability of this contact-based micromechanical technique to differentiate natural EV populations based on stiffness.^[9–14]^ One study investigated the relationship between exosome stiffness and exosome interactions with an endothelial layer.^[11]^ Compared to exosomes from non-malignant cell lines, malignant cell-derived exosomes were significantly softer and less adhesive, which likely contributed to the observed increased endothelial monolayer disruption and transendothelial transport of the malignant exosomes. This exosome-endothelial layer relationship may increase tumor cell extravasation and metastasis. Another study reported that EV stiffness correlated with breast cancer cell line malignant transformation.^[15]^ The stiffness, specifically linear elastic Young’s modulus, decreased as cells progressed from a non-tumor state to noninvasive to invasive phenotypes. When assessing the physical properties of submicron EVs derived from breast cancer patient plasma in the same study, the authors measured higher concentration, wider size distribution, and softer EV counts for the more dangerous invasive and ductal in situ carcinoma compared to benign samples. A comparison of hepatic cell lines indicated that EVs from different origins possess different lipid compositions that relate to the origin cell and exhibit different mechanical properties.^[13]^ These studies and others indicated that EV stiffness could be a clinically useful intrinsic biomarker. EV mechanical properties are also worthy of continued study for the fundamental understanding of diseases. For the blood disorder hereditary spherocytosis, red blood cells release more EVs (vesiculation) compared to healthy conditions.^[16]^ An AFM study revealed a possible mechanism for this increased rate of vesiculation: red blood cell EVs from hereditary spherocytosis patients had altered protein content that softened the EV membrane, which could make EV budding easier.^[12]^

However, AFM’s technical challenges will limit the field’s ability to fully understand EV stiffness implications in disease mechanisms and its potential use as a clinical biomarker. The AFM strengths of high spatial resolution and sensitive force control and detection are offset by the technique’s low throughput. AFM measurements are relatively slow, only 1 – 20 cells per hour depending on the AFM operation mode and level of detail needed to measure the desired elastic or viscoelastic mechanical property.^[17]^ AFM low throughput makes it challenging to measure a large number of EVs that is needed to achieve good statistics and capture the complexity of inherently heterogeneous biological EVs.^[18,19]^ A recent effort has demonstrated an AFM procedure that simultaneously captures EV morphology information and indirectly estimates mechanical properties from the contact angle of EV with a substrate of known surface properties at a rate of several hundred nanoscale EVs per hour.^[20]^ Despite yielding only semiquantitative mechanical properties, this higher throughput AFM approach was able to discriminate between vesicles of similar size but different stiffnesses.

Another limitation is the lack of standardized approaches for AFM mechanical measurements. Currently, there is large variability between reported mechanical EV measurements by AFM across groups from Young’s moduli of approximately 1 – 1500 MPa.^[11,15,19,21,22]^ These EV stiffness values are suspiciously high given the reported stiffness measurements of giant plasma membrane vesicles harvested from cells. Giant plasma membrane vesicles, generated by inducing membrane blebbing of cells, are very soft in fluid environments with low bending stiffnesses.^[23]^ Major factors contributing to this variability and suspiciously high stiffnesses are the application of different mechanical models to interpret stiffnesses from the AFM force – indentation curves and differences in experimental procedures. Recently Vorselen et al. proposed a standard operating procedure for EV stiffness measurements by AFM.^[18]^ Their proposed mechanical model accounts for the pressure build-up inside the vesicle due to substrate adhesion, which is often neglected in AFM EV studies^[24]^ and could explain the reported mega-Pascal moduli. Studies with this approach appear more consistent across sample types and plausible stiffness values. For example, trends of nanoscale EV stiffness with EV size and lamellarity (number of lipid bilayers) were well captured.^[24,25]^ The AFM mechanical modeling and experimental protocol challenges arise mostly due to AFM’s contact-based deformation mode. When squeezing a soft nanoscale object between a hard AFM probe and hard substrate, it is easy to enter a non-linear deformation regime, possibly violating the chosen model’s assumptions if the operator is not diligent and careful. The osmotic pressure effects that appear to contribute to effective stiffnesses measured by AFM are hard to measure independently. These pressures cannot be separately measured during AFM indentation of EVs for stiffness and must be estimated from separate experiments.^[24]^ Other techniques based on non-contact deformation modes could have a simpler deforming force that makes for easier and more accurate mechanical modeling for extracting EV stiffness from the observed deformation.

All the previous studies mentioned so far have exploited AFM’s nanoscale spatial resolution to study nanoscale extracellular vesicles less than 1 μm in diameter. This is reasonable given the importance of submicron EVs in physiology and pathophysiology and the fact that AFM is the only technique with the capability to measure nanoscale mechanics.^[9]^ However, one may wonder about the unusually large extracellular vesicles (L-EVs) > 1 μm that have not been studied to the same extent. What information do these EVs contain that has not been explored? L-EVs could be easier to handle and separate due to their large size and the ability to see them easily with conventional optical microscopy. The L-EVs’ volumes likely mean more biomolecular cargo that could make assays more accurate and robust for (large) EV subpopulation characterization compared to exosomes in many instances. The relatively few L-EV studies demonstrate that they are compositionally distinct from smaller exosomes and can activate specific intercellular communication pathways. Oncosomes, L-EVs that contain oncogenes, contain different protein cargo compared to exosomes.^[26]^ A comparative study of large (1-10 μm) and small (1-200 nm) EVs from prostate cancer cells showed that the large EVs contained active kinases that activated fibroblasts to promote tumor growth while smaller exosomes elicited a much smaller response.^[27]^ Therefore, studies of microscale L-EVs are worthy to pursue and also appropriate for microscale techniques that are unsuitable for nanoscale EVs.

We recently reported non-contact stiffness measurements of EVs using microfluidics.^[28]^ We adapted the approach from our biomechanical studies of suspended cells^[29]^ and synthetic vesicles^[30]^ to measure the stiffnesses of spherical L-EVs (1-5 μm diameter) isolated from human blood plasma. EVs were gently stretched in planar extensional flow, like a traditional tensile test from engineering. By measuring L-EVs in a natural fluid environment and low-force regime without hard probes and surfaces, we achieved a closer agreement with linear mechanical theory and therefore more accurate stiffness measurements. We measured L-EVs to have lower effective stiffnesses than suspended cells (~200 times softer), which is more reasonable given that EVs lack the relatively stiff cell nucleus and cytoskeleton. Though feasibility was demonstrated, the limits of our technique’s detection sensitivity need to be established. In this work, we tested if our microfluidic micromechanical approach could discriminate between cell lines and detect changes in L-EV biomolecular composition.

The altered EV biomolecular composition test case for this work was a genetic mutation common in glioblastoma multiforme (GBM) brain cancers, known to affect lipid metabolism. Mutations in isocitrate dehydrogenase 1 (IDH1) occur in many GBM brain cancers^[31]^ and has also been linked to several other cancers. Evidence suggests that this IDH1 mutation could be an early driver of oncogenesis, but the specific mechanisms and contributions remain controversial.^[32]^ The IDH1 mutation’s oncogenesis contributions could include a lipid metabolic disorder,^[33]^ leading to possible changes in lipid content and rearrangements of lipid order in the cell plasma membrane and membranes of internal organelles.^[34]^ Overall, knowledge of the changes to lipid composition and structure could lead to a more complete understanding of the consequences of IDH1 mutations on cellular functions and ultimately the discovery of new therapeutic targets.

EV stiffness can be a useful metric to capture the expected lipid changes due to the IDH1 mutation. Tightly packed biomolecules result in stiffer biomembranes compared to loosely packed biomolecules.^[35]^ Specifically, the change in the degree of lipid saturation potentially manifests as membrane stiffness changes. Unsaturated lipids that contain double carbon bonds will pack less tightly than saturated lipids,^[36]^ leading to a more fluid, or less stiff, biomembrane.^[37]^ Therefore, stiffness could quantify the physical effect of IDH1 lipid dysfunction on membrane biomechanics. Characterizing the variation of stiffness of EV with IDH1 mutation is useful to understand the biophysical ramifications of lipid disruption.

Here, we characterized the stiffness of large EVs from several GBM cell lines and investigated the IDH1 mutation’s effect on the EVs’ phospholipids, proteins, and stiffness. We use very general terminology ‘large extracellular vesicles’ (L-EVs) to refer to the atypically large EVs (>1 μm) derived from cancer cells that were studied here. L-EVs were isolated from the GBM cell-line culture supernatants using centrifugation. We used our non-contact microfluidic platform to measure the biomechanical properties of individual L-EVs (**Figure 1**). We first measured the stiffnesses of L-EVs from different GBM cell lines to determine if there were detectible differences in effective L-EV stiffnesses across cell lines within the same disease model. In some cases, L-EV stiffness from different cell lines displayed statistically distinct distributions. Next, we tested a hypothesis that the IDH1 mutation known to affect lipid metabolism could be detected as changes in L-EV stiffness. We considered two types of GBM cell lines, MGG and Gli36, and compared the stiffnesses of L-EVs from wild-type and IDH1 mutant cells. Complementary lipidomic analysis and protein content measurements were used to interpret the results from microfluidic stiffness measurements that IDH1 mutant cell line-derived L-EVs were stiffer than those from the wild-type counterparts.

**Figure 1.**
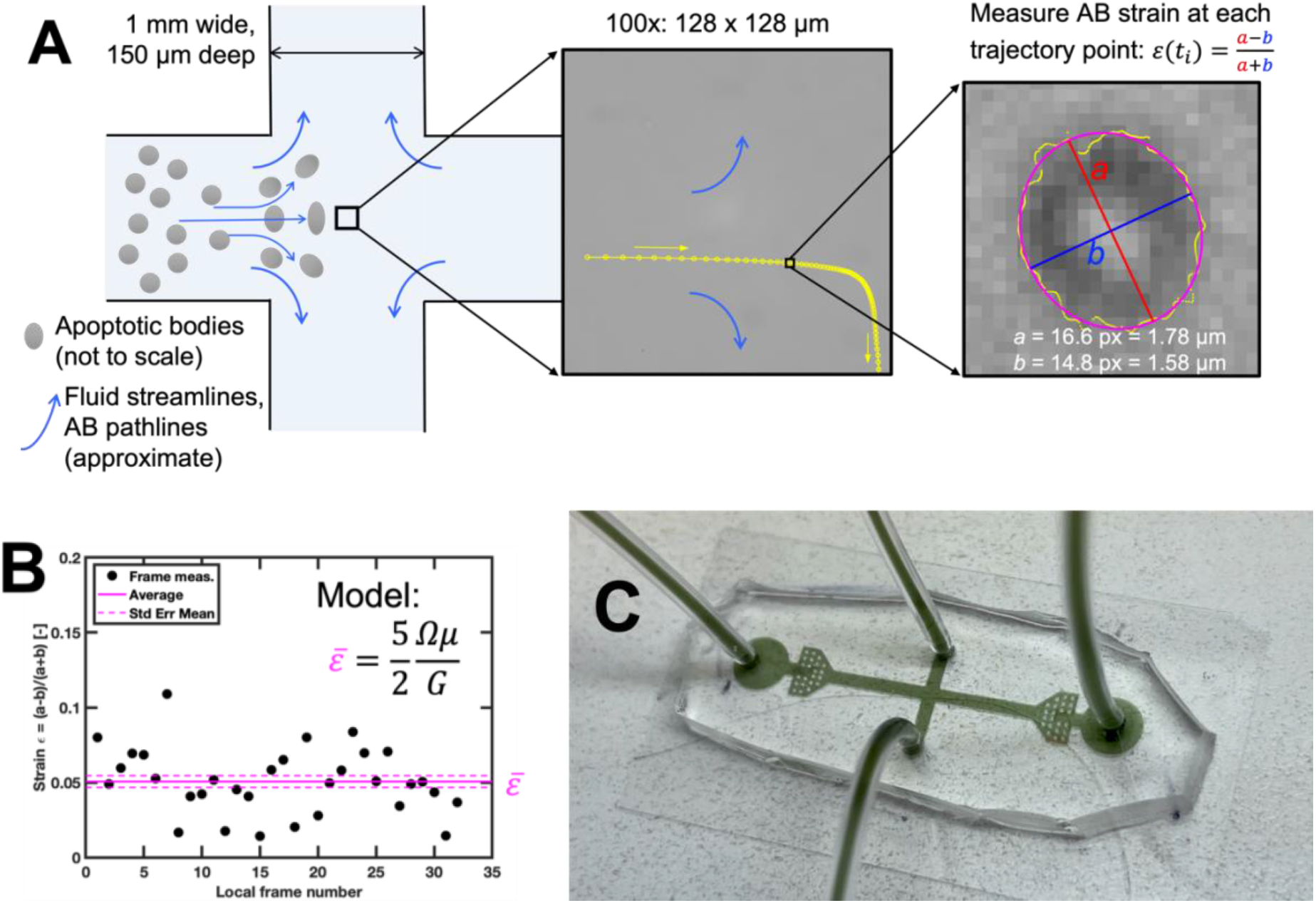
Non-contact microfluidic technique to measure the effective stiffness of microscale large extracellular vesicles (L-EVs, diameters >1 μm). **(A)** A schematic of the strain measurement procedure. Suspended L-EVs were gently elongated by extensional flow at the center of a cross-slot microfluidic device. At each time point along the L-EV’s trajectory, the L-EV stretch was quantified from a strain measure based on the axes of the best-fit ellipse to the detected contour. **(B)** Extracting an effective stiffness (*ε*) from the L-EV’s average stretch. A linear elastic model assumed that the L-EV is an isotropic linear elastic sphere that is deformed in planar extensional flow. The model was applied to extract the effective shear modulus *G* from the average strain. *Ω* is the constant extensional strain rate of the suspending fluid at the center of the extensional flow microfluidic device. *μ* is the viscosity of the suspending fluid (0.6% and 0.7% w/v methyl cellulose in PBS). **(C)** Extensional flow microfluidic device that generated planar extensional flow with a constant extensional shear rate to deform L-EVs.

## 2. Results

### 2.1. Large extracellular vesicle population characterization

L-EV morphology was observed within the microfluidic device (**Figure 2**). Many L-EVs were approximately spherical with visibly uniform appearance and composition, though some were irregular clumps (**Figure 2A**). Only spherical L-EVs were measured for effective stiffness because the spherical shape and homogenous composition, at least the microscale as visible in optical microscopy, indicated that these L-EVs could be reasonably approximated as isotropic, homogenous linearly elastic spheres in the mechanical model. The diameter of all L-EVs measured for stiffness ranged from 0.65 – 5.7 μm with a sample mean of 2.6 μm and sample standard deviation of 0.95 μm (**Figure 2B**). The range of L-EVs measured here was within the 1 – 5 μm size range considered for atypically large vesicles. The disaggregated size distributions from all L-EV populations measured for stiffness are presented in Figure S1 the Supporting Information. Estimates of L-EV population hydraulic diameter (D_H_) were measured using dynamic light scattering (DLS). The L-EV size distributions from DLS (D_H_ = 3.2 ± 1.2 μm to 4.2 ± 1.6 μm, mean ± standard deviation) (**Figure S2**) matched the expected size range for L-EVs (1 – 5 μm) and were consistent with the size distribution of L-EVs measured for stiffness (**Figure 2B**).

**Figure 2.**
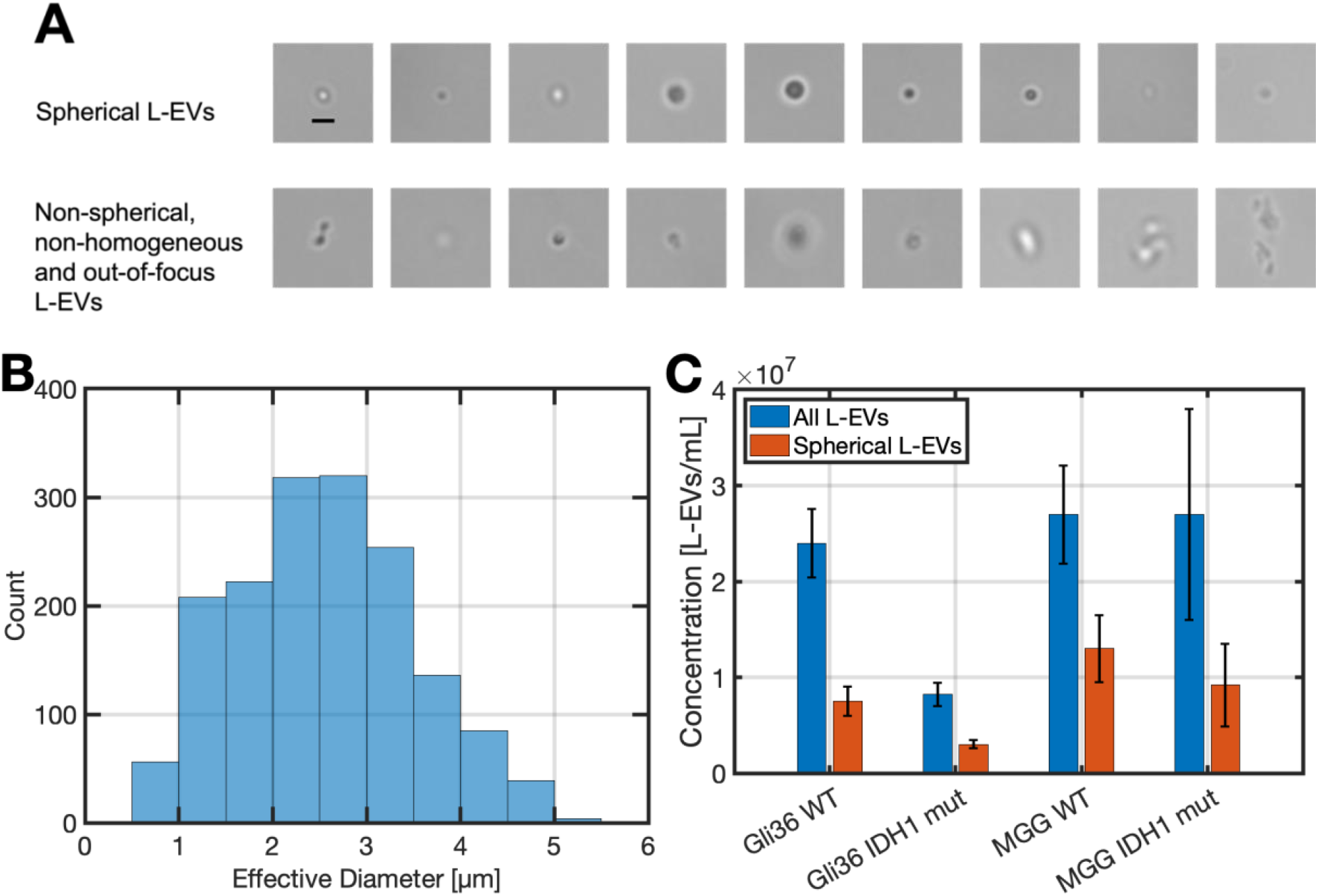
Characterizations of large extracellular vesicle (L-EV, diameters >1 μm) morphology, size, and concentration. **(A)** Suspended single L-EVs being gently stretched by extensional fluid flow in the extensional flow region of the microfluidic device. Spherical, homogeneous L-EVs (top row) that matched the assumptions of the mechanical model (isotropic linear elastic solid material, spherical initial shape) were measured for effective stiffness. L-EVs that were out of focus or had a far from the spherical shape (bottom row) were not considered for stiffness measurements because they violated the model’s assumptions. Larger aggregates were also observed in fluid chambers compared to these observations within the microfluidic device. **(B)** Distribution of the effective diameters of all spherical L-EVs measured for stiffness (*N* = 1643). **(C)** Concentration estimates from representative L-EV populations from MGG and Gli36 wild-type and IDH1 mutant cell lines. Concentrations were estimated from z-stack images captured within the microfluidic device when the fluid was not moving. Bar heights indicate the average of 4 – 13 z-stack measurements in different locations. Error bars are the standard error of the mean of these measurements.

The concentration of all suspended biological material was roughly uniform across representative samples (**Figure 2C**). Within the microfluidic device, the concentration of all material was 10 – 30 million objects per milliliter. The concentration of spherical L-EVs that were measured for stiffness was a little less than half of the total, or roughly 0.4 − 1.2 × 10^7^ spherical L-EVs per milliliter.

Qualitatively, we observed larger aggregates within the suspended L-EV samples when imaging in a droplet or fluid chamber compared to observations within the microfluidic device. After defrosted L-EVs pellets were resuspended in aqueous methyl cellulose in phosphate buffered saline (PBS), the solution was pipetted into fluid chambers or droplets on a glass slide. Large aggregates greater than ~10 μm in size were common while isolated <5 μm spherical L-EVs were difficult to find (**Figure S3**). However, within microfluidic devices these same L-EV suspensions were distinctly different with many isolated <5 μm spherical L-EVs and smaller aggregate clumps <10 μm in size. During infusion into the microfluidic device, the suspended L-EVs and aggregates were presumably broken up by the fluid shearing forces experienced in the syringe, needle, tubing, and device entrance hole. Microfluidics is a gentle micromechanical technique compared to contact-based AFM and micropipette aspiration. However, our observations indicate that biological materials were subject to fluid shearing forces that probably broke up large, loosely bound aggregates.

### 2.2. Large extracellular vesicle stiffness varies between some GBM cell lines

L-EVs from different GBM cell lines had varying stiffness distributions as measured by the extensional microfluidic technique (**Figure 3**). According to the two-sided Wilcoxon rank sum test, some L-EVs stiffness distributions come from populations with different means for p-values 0.05 or lower. The degree of statistical significance varied with some cell lines producing L-EVs that were more distinct stiffness-wise than other cell lines. For example, T98G L-EVs were significantly softer than the U87- and MGG-derived L-EVs (p < 0.01). L-EVs from Gli36, U87, and MGG cells were not statistically significantly different according to the p < 0.05 threshold in pairwise comparisons with the two-sided Wilcoxon rank sum test. A multi-group comparison with the nonparametric Kruskal-Wallis test drew a similar conclusion: the Gli36, U87, and MGG L-EVs stiffness measurements came from the same population or different populations with the same mean. A summary of the statistical analyses is provided in the Supporting Information **(Table S1)**. These data indicate that L-EV stiffness distributions could potentially be used to help assess cellular origins with our microfluidic approach following further development to improve measurement accuracy and sensitivity.

**Figure 3.**
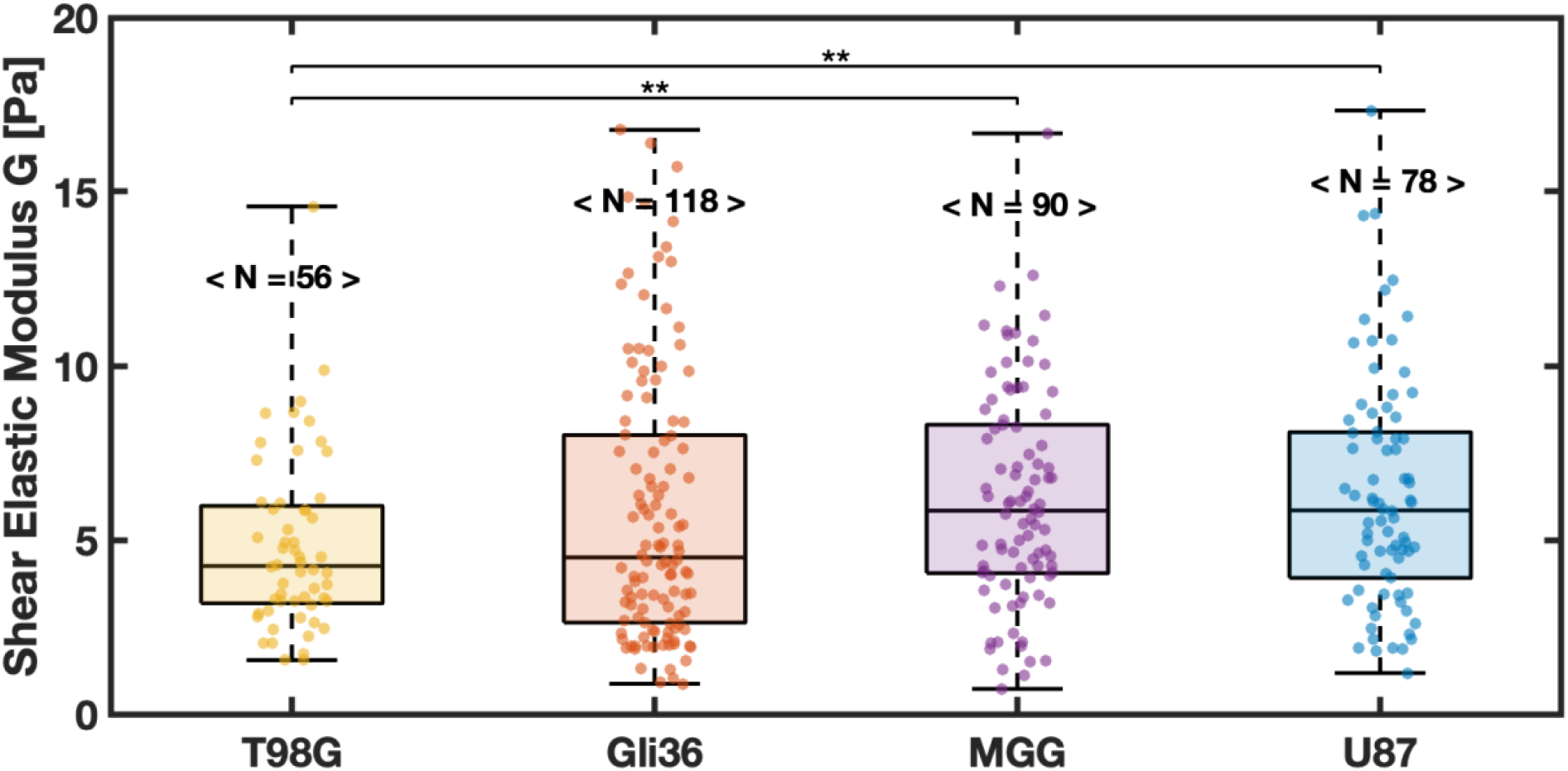
Stiffness distributions of large extracellular vesicles (L-EVs, diameters >1 μm) from different glioblastoma multiforme (GBM) cell lines. All measurements were performed with L-EVs suspended in 0.6% methyl cellulose in phosphate buffered saline (PBS, viscosity *μ* ~ 30 mPa·s). The bracketed numbers indicate the number of individual L-EV stiffness measurements per distribution. Box plots: middle line is the median, box top and bottom edges are the 25^th^ and 75^th^ percentiles, whiskers indicate the range (minimum to maximum) of the distribution, and individual L-EV stiffness measurements are indicated by the scatter plot overlayed on the box plot. Statistical analyses of pairwise comparisons were conducted with the two-sided Wilcoxson rank sum test with statistically significant p-values shown as *p < 0.05, **p < 0.01.

All L-EVs measured had shear elastic moduli within *G* = 1 – 20 Pa (equivalent to Young’s moduli *E* = 3*G* = 3 – 60 Pa for incompressible materials, as is assumed in our mechanical model). We measured L-EVs stiffnesses to be softer than suspended whole cells, whose stiffnesses range from *E* = 0.1 – 100 kPa as measured by various techniques, including non-contact microfluidic approaches.^[38]^ This is reasonable given that the cell nucleus and cytoskeleton have been shown to often dominate overall cell effective stiffness.^[39]^ L-EVs are composed of a heterogeneous lipid bilayer membrane with various proteins crossing or adhered to the bilayer, which can be expected to be softer than the nucleus and cytoskeleton. The results here closely agree with the stiffnesses for L-EVs derived from blood plasma reported in our previous work.^[28]^ The plasma L-EV shear moduli ranged from 1.2 Pa to 14.2 Pa with an average of *G* = 5.6 ± 0.5 Pa (mean ± standard error of the mean, *N* = 50). The consistency in L-EV shear elastic moduli between our two studies suggests that the dynamic range of L-EV stiffness across cell lines and cell types, at least for mammalian cells, is small, and therefore measurement techniques should be very sensitive.

All stiffness distributions except MGG are non-normal according to the Lilliefors test (5% significance level) (**Figure 3**). This likely arises because strains are small and in the linear regime (strain < 0.1) and close to the natural limit of zero strain or no stretch.

### 2.3. Increase of the large extracellular vesicle stiffness with IDH1 mutation

We next investigated the stiffness of L-EVs from wild-type and IDH1 mutation cell lines expressing the IDH1 mutation protein (**Figure S5**). In two replicate measurements of each cell line, L-EVs from GBM cells with the IDH1 mutation showed higher stiffness compared to those from wild-type GBM cells according to the Wilcoxon rank-sum test with p-values < 0.05 in all cases (**Figure 4**). Essentially, the measured stiffness differences for the wild-type vs. IDH1 mutation within a cell line (**Figure 4**) were more distinct and had smaller p-values than the comparisons across GBM cell lines (**Figure 3**). This may indicate that the IDH1 mutation causes more alterations to the cell membrane in terms of biomolecular content and arrangements than is common across GBM disease model cell lines.

**Figure 4.**
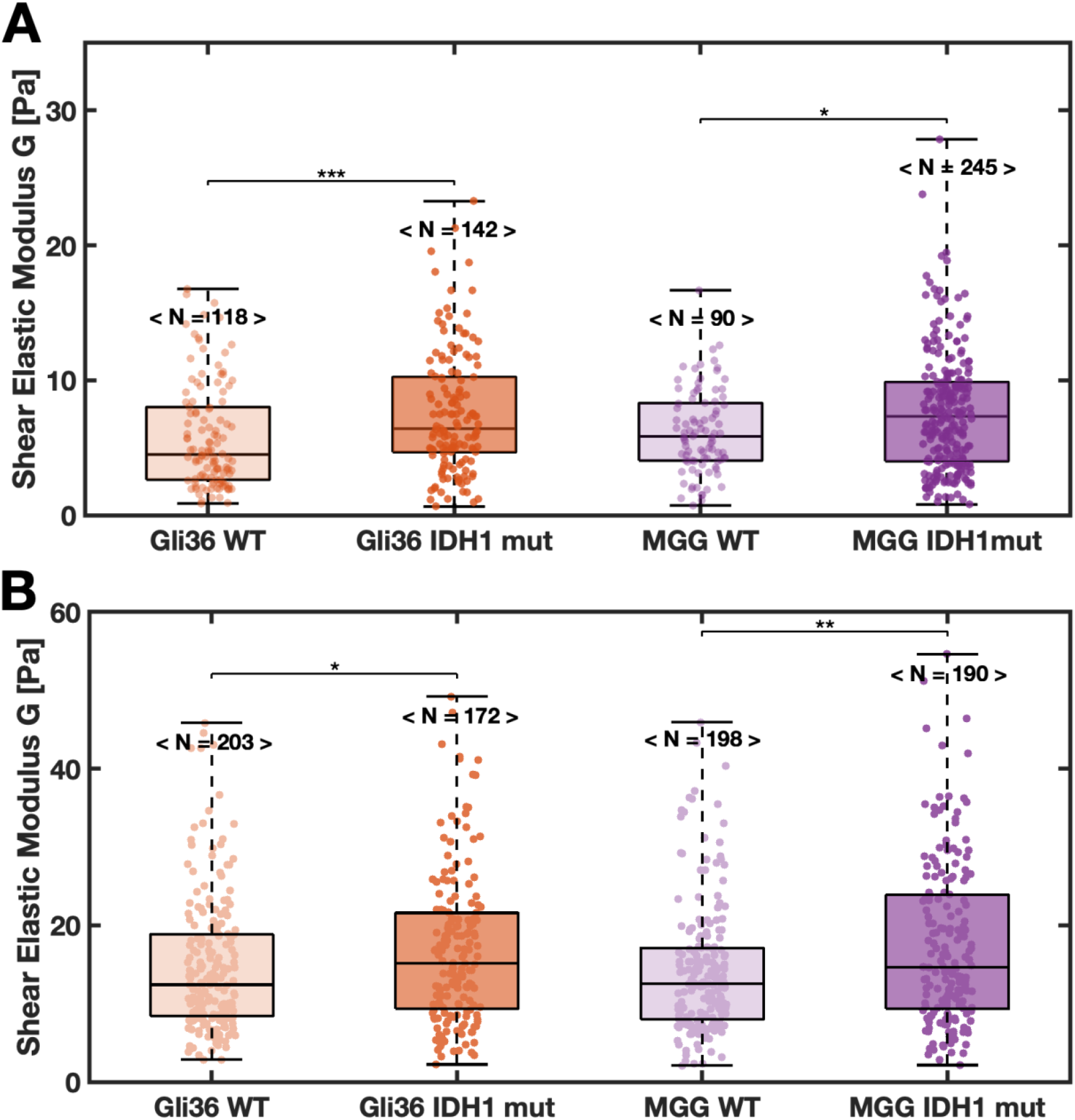
Large extracellular vesicles (L-EV, diameters >1 μm) from two different glioblastoma multiforme (GBM) cell lines become stiffer with the IDH1 mutation. Stiffness distributions of L-EVs derived from Gli36 and MGG GBM cell lines as measured in two independent experiments. The dimensions of the microfluidic devices remained the same for both sets of experiments, meaning similar extensional strain rates. The suspending solutions with different viscosities were used in the two independent experiments. **(A)**Gli36 wild-type and IDH1 mutant cell lines and MGG wild-type and IDH1 mutant cell lines in the first experiments performed in 0.6% methyl cellulose in phosphate buffered saline (PBS, viscosity *μ* ~ 30 mPa·s). **(B)**The second experiments performed in 0.7% methyl cellulose in PBS (viscosity Gli36: *μ* = 65 mPa·s, MGG: *μ* = 68 mPa·s). The bracketed numbers indicate the number of individual L-EV stiffness measurements per distribution. Box plots: middle line is the median, box top and bottom edges are the 25^th^ and 75^th^ percentiles, whiskers indicate range (minimum to maximum) of the distribution, and individual L-EV stiffness measurements are given by the scatter plot overlayed on the box plot. Statistical analysis of pairwise comparisons were conducted with the two-sided Wilcoxson rank sum test with statistically significant p-values shown as *p < 0.05, **p < 0.01, ***p < 0.001.

To test the repeatability and accuracy of our microfluidic measurement technique, a higher viscosity suspending solution was used in the second independent experiment. If the mechanical model captures the major biomechanical phenomena of the L-EV deforming in the suspending fluid, then the controllable experimental parameters can be changed and the same EV stiffness would still be measured. For our technique, the controllable experimental parameters are the suspending fluid viscosity *μ* and the uniform fluid extensional strain rate *Ω* at the center of the cross-slot microfluidic device. Fluid viscosity was determined by the concentration of methyl cellulose dissolved in PBS. For the first independent experiment (**Figure 4A**), the suspending fluid was 0.6% w/v methyl cellulose in PBS. For the second independent experiment (**Figure 4B**), the suspending fluid was 0.7% w/v methyl cellulose in PBS. The remainder of the experimental control parameters—device dimensions, height of the imaging focal plane above the device’s glass bottom, and fluid flow rates—were the same between the two independent experiments. For the pairwise comparisons within an independent experiment, care was taken to make all experimental control parameters as identical as possible by using the same device used and the same batch of suspending fluid for the wildtype and IDH1 mutant cases. As seen when comparing Figure 4A and 4B, the EV stiffnesses extracted by the mechanical model were slightly different from the experiments in the higher viscosity 0.7% w/v methyl cellulose in PBS leading to ~3 times higher absolute stiffness values. For instance, in 0.6% w/v methyl cellulose wild-type Gli36-drived L-EVs had a stiffness of 5.8 ± 0.4 Pa (mean ± standard error of the mean) while in 0.7% w/v the stiffness was measured to be 14.7 ± 0 .6 Pa. However, the statistical trends were consistent: When experimental parameters were held constant within the two independent experiments, wild-type Gli36 and MGG-derived L-EV stiffness distributions were stiffer than the wildtype counterparts.

We attribute the main source of the discrepancies in absolute stiffness values between Figure 4A and 4B to the uncertainty in fluid viscosity. The other experimental parameters—device dimensions, flow rates used to stretch L-EVs, and the imaging plane location within the device—remained practically constant; only the fluid viscosity was significantly changed. The mechanical model assumes that the fluid is a Newtonian fluid, meaning the viscosity is constant no matter how fast the fluid is flowing. In reality, aqueous methyl cellulose in PBS is slightly shear-thinning as measured by us **(Figure S4)** and other groups.^[40]^ The precision of our microfluidic L-EV stiffness measurements could be improved by using a suspending fluid with a more constant viscosity (closer to ideal Newtonian behavior). Then changing the suspending fluid viscosity would yield more similar stiffness values. Despite this, the result of stiffer L-EV with IDH1 mutation in Figure 4 was consistent within independent experiments in which experimental parameters were constant, lending confidence that this observed trend is due to the IDH1 mutation.

### 2.4. Altered lipid profile of large extracellular vesicles derived from cells with the IDH1 mutation

To understand the stiffness increase in L-EVs from GBM cells with IDH1 mutation, we conducted lipidomics analyses on four L-EV samples (L-EVs from GLI36 and MGG cell lines in wild-type and IDH1 mutation, respectively; **Figure 5**). The numbers of detected lipids in positive ion species and negative ones were 264 and 90, respectively. We conducted principal component analysis (PCA) to assess the statistical significance of the lipid difference between samples (**Figure S6**). In the PCA analysis, the analyzed samples were in different quadrants, indicating that the lipid components of each sample showed distinct characteristics. We first analyzed the proportion of different lipid types and their differences between wild-type and IDH1 mutation cell lines (**Figure 5A** **and** **5B**). It shows the amount of phospholipids, a major component of membranes, decreased in both GLI36 and MGG cells with the IDH1 mutation. Among phospholipids, phosphatidic acid, phosphatidylcholine, and phosphatidylethanolamine were decreased in the L-EVs derived from IDH1-mutated cells (**Figure 5C, 5D** **and S7**). In particular, a large number of unsaturated lipids was reduced by a greater portion than saturated lipids while the numbers of increased lipids were the same between saturated and unsaturated lipids (**Figure 5E** **and** **5F**). This is evidence that for these GBM cell lines, the IDH1 mutation reduced the amount of unsaturated lipids. Compared to the L-EVs derived from the wild-type cells, the IDH1 mutated L-EVs had more saturated lipids.

**Figure 5.**
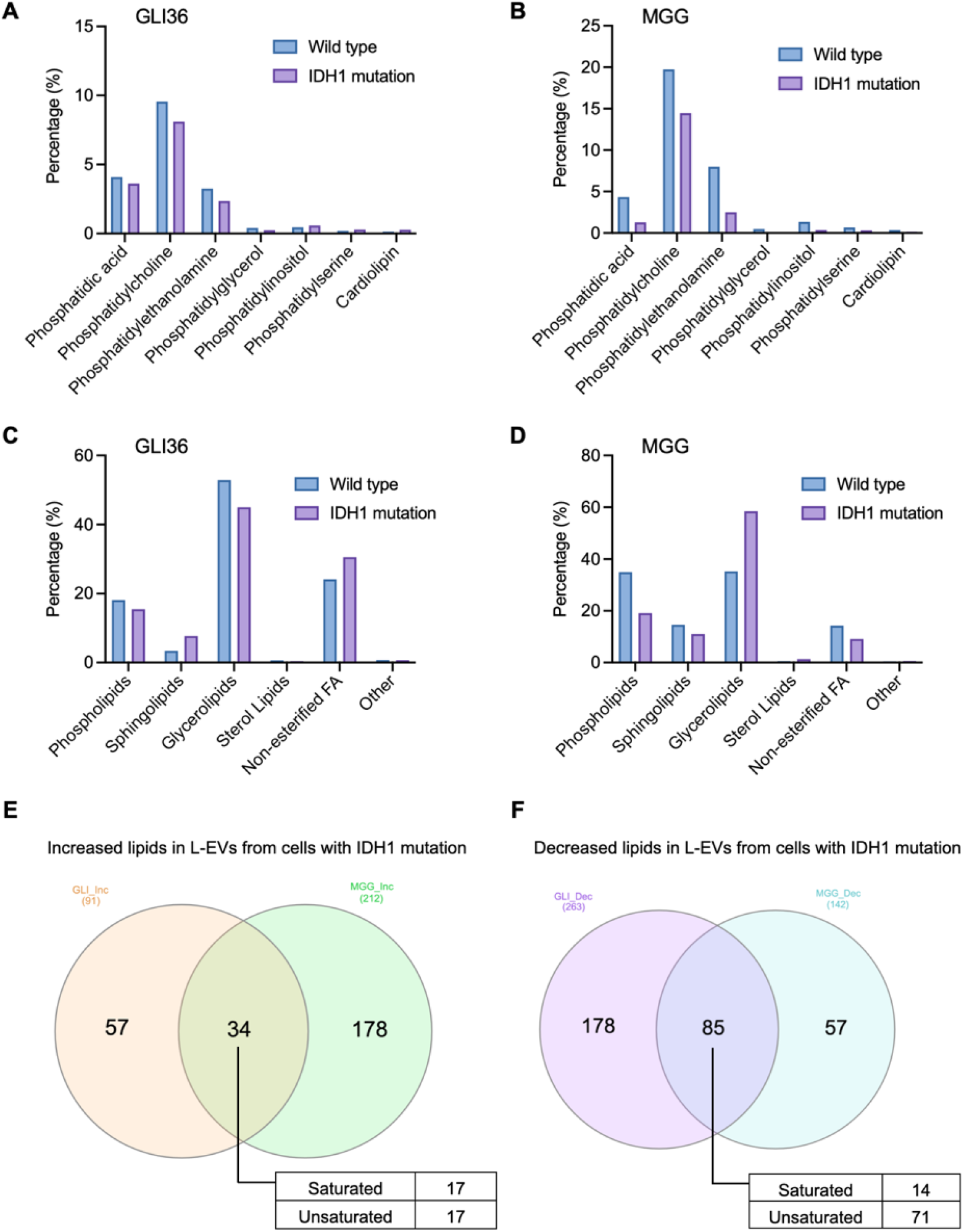
Lipid profiling comparison between large extracellular vesicles (L-EVs, diameter >1 μm) derived from wild-type and IDH1-mutated glioblastoma multiforme cell lines (MGG and Gli36). (**A-B**) The percentage composition of lipid categories in L-EVs from Gli36 (A) and MGG (B) cells normalized by the total lipid amounts. (**C-D**) The percentage composition of phospholipid categories in L-EVs from Gli36 (C) and MGG (D) cells. (**E-F**) Venn diagrams of the numbers of increased or decreased lipids in L-EVs from IDH1-mutated Gli36 and MGG cells relative to L-EVs from the wild-type cells. Equal numbers of saturated and unsaturated lipids were increased in L-EVs from the IDH1-mutated cell lines (E). In contrast, a larger number of unsaturated lipids was decreased in L-EVs from IDH1-mutated cell lines (F).

## 3. Discussion

Here, we demonstrated that our non-contact microfluidic approach to measuring L-EV effective stiffness, specifically the shear elastic modulus *G*, is sufficiently sensitive in some cases to detect differences due to mutations and across cell lines. Our previous work demonstrated that the microfluidic technique is a gentler alternative to atomic force microscopy for L-EVs and yields reasonable order-of-magnitude measurements of L-EV stiffness.^[28]^ The present work challenged the resolution limits of our approach to discriminate between GBM cell lines and between wildtype and IDH1 mutant cell populations. This technique appears well-suited for instances in which contrasting biomembrane composition is expected. For the IDH1 mutation that disrupts lipid metabolism, all independent experiments comparing L-EV stiffness distributions between wildtype and IDH1 mutated cells demonstrated statistically significant differences (p < 0.05 or better). L-EV stiffness distributions from GBM cell lines were less distinct. Only T98G-derived L-EVs were different from MGG- and U87-derived L-EVs with p < 0.05. However, if a lower threshold for statistical significance were accepted (p < 0.1), then the Gli36 L-EVs could be considered softer compared to the MGG and U87 L-EVs. The technique’s L-EV subpopulation identification capability might be improved through more accurate determinations of the suspending fluid’s viscosity and extensional strain rate.

The results point at the possibility of using L-EV stiffness as non-invasive glioma biomarker to detect lipid changes due to mutations or other altered cellular processes. The lipidomics data indicated that the IDH1 mutant L-EVs were stiffer at least in part due to a relatively higher amount of saturated lipid content compared to those derived from the wildtype cells. In general, a biomembrane composed of saturated lipids is stiffer than one made of unsaturated lipids because saturated molecules can pack together more tightly, leading to larger attractive intermolecular forces between lipid fatty acyl chains and reduced membrane flexibility and increased rigidity.^[35]^ The microfluidic measurement technique appears capable of detecting this change in lipid composition through statistically significant differences in L-EV stiffness distributions between populations from wild-type and IDH1 mutant cell lines. Taken together, our evidence implied that the IDH1 mutation altered the lipid composition of GBM cell lines, specifically increasing the relative amount of the tightly packing saturated lipids, and that this change in lipid composition could be a driving cause of L-EV stiffening with the IDH1 mutation. By isolating GBM-derived L-EVs from patient blood samples, it may be possible to infer the presence of the IDH1 mutation or other biomembrane disrupting mutations, or lack thereof, by measuring the stiffness of L-EVs measured at high throughput with a microfluidic device like our technique.

In comparison to a previous study of the IDH1 mutation’s effects on lipids in organelles, our study of L-EVs also demonstrated that the IDH1 mutation results in measurable changes to lipid composition and downstream biophysical effects. When investigating the changes in lipid profiles in the organelles of the U251 GBM cell line using Raman spectroscopy, Lita et al. found that there were relatively more saturated lipids in Golgi apparatus for two versions of the IDH1 mutation (R132C and R132H).^[34]^ Their evidence indicated that IDH1 mutations increased the degree of lipid unsaturation and produced more monounsaturated fatty acids in the endoplasmic reticulum and Golgi apparatus. This led to morphological defects (swelling) in these organelles. We investigated L-EVs, which presumably bleb off from the outer cell plasma membrane, as opposed to internal cellular organelles. Given that lipids make up a large fraction of EV biomolecular content, we hypothesized that the IDH1 mutation effect of dysfunctional phospholipids is also present in EVs. For the IDH1 mutated MGG and Gli36 GBM cell lines studied here, the lipid profiles of their L-EVs demonstrated higher lipid saturated levels for the mutated case compared to the wild-type. The change in lipid saturation levels was consistent with the observed stiffening of these vesicles: IDH1 mutant L-EVs were significantly stiffer than the wild-type L-EVs in all independent experiments using two different GBM cell lines. Between the two studies, one could conjecture that the IDH1 mutation affects lipid composition to the point of changing the biophysical characteristics (morphology, stiffness) of small membrane-bound subcellular units both inside and outside the cells.

As with many biological measurements, there is a high degree of overlap in the stiffness distributions for L-EVs derived from different cell line groups. Thus, individual L-EVs cannot be classified to a GBM cell line based on stiffness alone, according to our results. However, hypothesis testing indicates that these L-EV groups come from different populations. Machine learning algorithms could potentially be used on these stiffness data along with other L-EV shape metrics (area, perimeter, circularity, strain or stretch, etc.) to classify and predict the cell line or L-EV population from which single L-EVs come from. Such classification studies based on whole cell deformation and shape have been done by others.^[41–43]^ These studies demonstrated that analyzing multiple physical phenotype metrics together can be more accurate at classifying cancer cell lines than a single metric (e.g., stiffness) alone.

The advantages of high-throughput and non-contact EV deformation mode in microfluidics are offset by a size limitation. We are limited to studying microscale L-EVs because our microfluidic technique requires visual measurements of single L-EV deformation from optical microscopy, which is limited by the diffraction of visible light. To improve the size resolution of microfluidics, other imaging modalities (e.g., digital holography) that have a smaller diffraction limit must be used. It may be possible to correlate changes to molecular spectra to nanoscale EV deformation or mechanical properties. Currently, AFM is the only mechanical measurement technique for nanoscale EV mechanical properties, though at a relatively low throughput and involving contact-based deformations. In our view, microfluidics and AFM can both provide accurate and precise measurements of EV mechanical behaviors in a complementary manner. Both should be used to gain a more complete understanding of EV mechanics.

## 4. Conclusion

Microfluidics is a promising approach to characterizing the biophysical properties of microscale EVs that complement existing measurement technologies. We demonstrated the potential of non-contact microfluidics to discriminate between EV subpopulations. With the current state of our technique, EVs from some GBM cell lines could be distinguished based on L-EV stiffness distributions. In the investigation of the IDH1 mutation that is known to affect lipids, microfluidics-measured L-EV stiffnesses were much more distinct. The finding of consistently stiffer IDH1 mutant L-EVs was supported by lipidomics data. We conjectured that changes in lipid saturation level with the IDH1 mutation changed lipid packing, which then led to stiffer L-EVs compared to wildtype. Further refinement of experimental control parameters would increase measurement accuracy and could improve the microfluidic technique’s ability to classify and determine EV subpopulations.

This work contributes to understanding the range of EV physical properties, an important step for their use in clinical applications. Knowing physical properties of L-EVs is important for purification and separation processes in which unwanted objects are separated out of patient samples based on biophysical characteristics. Most EV research has focused on smaller exosomes and microvesicles. Our results demonstrate that the less-studied L-EVs are also rich multiplex biomarkers for bioanalysis that have potential for disease diagnostics and therapeutics.

## 5. Experimental Section

*L-EV Isolation*: GBM cell lines (T98G, U87, MGG, Gli36) were grown in Dulbecco’s minimum essential medium (DMEM, Hyclone) containing 10% fetal bovine serum (FBS, Gibco) at 37°C in 5% CO_2_. L-EVs were isolated from cell culture media collected using differential centrifugation. The aspirated media were centrifuged at 300*g* for 10 minutes followed by 2,000*g* for 20 minutes. The supernatant was removed. The remaining pellets were stored as 20 μL aliquots at −80°C. Aliquots were defrosted for 5 minutes before microfluidic and other experiments.

*Dynamic Light Scattering:* Dynamic Light Scattering (DLS) was used to measure the size distribution of L-EVs in aliquots using a Horiba NanoPartica SZ-100 Series Nanoparticle Analyzer with the Horiba NextGen Project SZ-100 software (version 2.20). Defrosted 20-μL aliquots were resuspended in 1 mL deionized water and pipetted into a disposable cuvette. Nine to ten DLS measurements were averaged to estimate the hydrodynamic diameter (D_H_) and polydispersity index (PDI) of the suspended L-EV samples. D_H_ and PDI results were reported as the average ± standard deviation. The large measured PDIs > 0.7 indicated that the L-EV populations had very broad size distributions and that DLS is not the most suitable technique to measure size distributions for these L-EVs.

*Microfluidic Device:* Custom microfluidic devices that generate extensional flow were fabricated using standard soft lithography techniques.^[44]^ The extensional flow device, also called a cross-slot, consists of two straight channels that intersect at a right angle. This basic device design has been previously used to mechanophenotype cell populations in patient biofluids,^[45,46]^ estimate the mechanical properties of suspended,^[29,47]^ and measure the dynamics of synthetic vesicles in extensional flow.^[30,48,49]^ This microfluidic device design has been shown to generate the intended planar extensional flow in the central region of the device by several groups,^[48,50]^ including our past work.^[29,30]^ Masters were fabricated from a dry-film photoresist (Riston GoldMaster GM140 photoresist polymer film, DuPont) and laminated on stainless steel wafers with a hot-roll laminator (Akiles ProLam Ultra Laminator) using previously described procedures.^[51]^ Polydimethylsiloxane (PDMS) devices were fabricated by mixing PDMS based and curing agent (Sylgard 184, Dow Corning) in a 10:1 ratio, pouring over the master, and curing. Devices were bonded to No. 1 coverslips (Fisher Scientific, 50×24 mm, thickness 0.13 – 0.17 mm) to allow for imaging within the device using an oil-immersion 100x objective. The dimensions of the channel’s rectangular cross-section (*w* = 1 mm wide and approximately *h* = 150 μm deep) were chosen to be very large relative to the size of the L-EVs (< 5 μm diameter). With this large size discrepancy, the flow field around individual L-EVs is close to the planar extensional flow (no z dependence of velocity) that is assumed by the analytical mechanical model used to extract L-EV stiffness from its observed stretching. Furthermore, the channel length from the inlet holes to the cross-slot’s stagnation point region (the intersection of the two straight channels) where the extensional flow is generated was sufficiently long (5 mm) to ensure that the fluid flow is fully developed by the time it enters region.

*Suspending Fluid:* For L-EV stiffness measurements in the microfluidic extensional flow device, the suspending fluid was 0.6% w/v or 0.7% w/v methyl cellulose (Thermo Scientific, molecular weight 454.513 g mol^−1^) in phosphate buffered saline (PBS, Invitrogen, 10 mM phosphate and 150 mM sodium chloride, pH 7.3 to 7.5). The viscosity of the suspending solution was measured on a TA Instruments Discovery Hybrid series rheometer with a cone-and-plate geometry (40 mm diameter). Viscosity was determined from a flow sweep protocol between 1 – 1000 s^−1^ strain rates with 10 points per decade. The viscosity used to in the analytical mechanical model was taken to be the average viscosity of 3 independent measurements. Within each independent trial, the viscosity was found by averaging the viscosity in the strain rate region after initial transients had died out and the normal stress was linear with shear rate. This averaging region corresponded to the shear regime starting from 5 – 10 s^−1^ up to 50 s^−1^. The average viscosity was approximately 30 mPa·s for the 0.6% w/v methyl cellulose in PBS and 65 – 69 mPa·s for the 0.7% w/v solution. Our results showed that methyl cellulose was slightly shear thinning (viscosity decreases with increasing shear rate), as observed by others.^[40]^ The L-EV mechanical property measurements were confined to a narrow range of extensional shear rates, ~2.8 – 8.3 s^−1^, and over this range we assumed that viscosity was approximately constant. This slightly non-Newtonian fluid behavior of the suspending fluid added uncertainty to our L-EV measurements, as noted in the Results.

*Microfluidic Experiments:* The details of our non-contact microfluidic technique that gently stretches individual microscale extracellular vesicles to measure their stiffnesses was detailed in our recent publication.^[28]^ Briefly, L-EV aliquots were defrosted for 5 minutes and then suspended in 700 μL of 0.6% or 0.7% w/v methyl cellulose in PBS (mcPBS) by gently pipetting to thoroughly mix the L-EVs throughout the solution. The extensional flow device was filled with mcPBS from both inlets to ensure no bubbles were present. Then the syringe containing the L-EV suspension was installed on one of the inlets. Fluid was infused into the extensional flow device from two opposite channel inlets (one side blank mcPBS fluid and the other the L-EV suspension in mcPBS) using a Harvard Apparatus PHD2000 syringe pump. The two inlet streams impinged on each other in the stagnation point region (the intersection of the two straight channels) to generate planar extensional flow. L-EV stretching in the stagnation point region was observed on a Zeiss Axiovert 200M inverted microscope using a 100x objective (working distance 0.13 mm). Movies were recorded with a Photometrics Prime95B sCMOS camera (9.346 pixels μm^−1^, full frame field of view 1200 by 1200 pixels or 128.4 μm by 128.4 μm, 16-bit images, 1 ms exposure time, ~40 frames per second) with the camera controlled by MicroManager.^[52]^ Fluid infusion flow rates were varied in increasing order from 500 μL hr^−1^ to 1500 μL hr^−1^ (equivalent strain rates approximately 2.8 – 8.3 s^−1^)^[29]^. There was at least 1 minute of system equilibration time was observed after increasing the flow rate prior to recording L-EV stretching movies at the higher flow rate. At each flow rate, 2 – 3 movies (500 frames recorded over 12.5 seconds) were captured at 35 μm focal plane height above the glass bottom and 2 – 3 movies at 40 μm focal plane height. The working distance limitation of the 100x objective limited movie capture focal planes a maximum of 35 - 40 μm above the device’s glass bottom, as opposed to the ideal location at the mid-channel height ~75 μm above the bottom due.

*Microscopy Image Analysis to Measure L-EV Deformation:* All image processing was conducted with a custom analysis code in MATLAB (MathWorks, R2020b) that used the MATLAB Image Processing Toolbox. First the smoothed background was subtracted from all movie frames. The background was the time-average image of all movie frames. It was smoothed using a Gaussian filter (9×9 pixel neighborhood, standard deviation of 2 pixels). The background-subtracted movie frames were binarized using a global threshold value that was manually selected prior to analysis to detect most microscale spherical L-EVs. Potential L-EVs were detected in movie frames using thresholding. Detected objects with areas (*A* = 0.5 – 10 μm^2^) and circularities (*C* = 4π*A*/*L*^2^ > 0.9) in the range expected for L-EVs were considered potential L-EVs while all other objects were discarded. The potential L-EV objects were linked into trajectories using the MATLAB adaptation^[53]^ of the particle tracking algorithm developed by John C. Crocker.^[54,55]^ All potential L-EVs in trajectories were visually checked for adherence to the L-EV mechanical analysis criteria of being in focus and close to spherical shape. Out-of-focus L-EV and clumpy aggregate trajectories were discarded from further analysis. Next, the edges (contours) of L-EVs along their trajectories were found. The background-subtracted movie frames were smoothed using a Gaussian filter (2×2 pixel neighborhood, standard deviation of 0.5 pixel). The L-EV contour location was defined as the location of maximum grayscale intensity gradient along rays (360 total) drawn from the center of the L-EV. Sub-pixel edge detection resolution was achieved by finding the maximum of a parabola fit to the local intensity gradient in a 5-pixel neighborhood. To obtain a consistent measurement of strain, an ellipse was fit to each L-EV contour using the least-squares method. The L-EV strain *ε* in a frame was defined as *ε* = (*a – b*)/(*a + b*) where *a* and *b* are the long and short axes of the fitted ellipse, respectively. The estimated strain for each L-EV was taken to be the average strain along its trajectory. L-EVs for which mean(*a – b*) < 1 pixel were discarded because this level of deformation was considered below the detection limit of the microscope system. A quality control step assessed the trends in the fitted ellipse area *A* and strain *ε* to judge edge detection quality and strain measurement certainty. Some L-EVs had scattered edge detection because the L-EV was rotating, it went out-of-focus along the trajectory, or the low image signal-to-noise level made contour detection highly variable. The following criteria filtered out L-EVs with high uncertainty in shape detection. If the range and scatter of either area and/or strain along the L-EV trajectory were too large and/or the number of frames in the L-EV trajectory was too small, then the average strain was considered a poor measurement and the L-EV was discarded from further analysis. Strain measurements were discarded if two or more of the following criteria for good edge detection were violated: std(*A*)/mean(*A*) < 0.25, *ε* range < 0.11, std(*ε*)/mean(*ε*) < 0.5, or number of frames in trajectory > 5.

*Analytical Mechanical Model to Extract L-EV Effective Stiffness:* L-EV stiffness was extracted from the L-EV strain measurements though analytical linear mechanical theory. This model treated L-EVs incompressible, isotropic linearly elastic solid spheres even though actual L-EV structure was more complicated with a complex membrane surrounding an aqueous core that contains biological cargo. Thus, in this study we measured the effective shear elastic modulus (effective stiffness) of L-EVs, i.e., the stiffness of an elastic sphere with similar mechanical behavior to the more complex L-EV in this microfluidic system. Due to the high objective power and relatively large channel dimensions, L-EV strain was measured close to the stagnation point and far from the channel walls (Figure 1A). At this location, L-EVs had experienced the extensional flow field for long enough time to be at their equilibrium stretch length. Therefore, our technique measured L-EV elastic (long-term) mechanical behavior. From the solution of an incompressible elastic sphere deforming in an arbitrary linear, low Reynolds number flow field,^[56]^ the following equation describes the sphere’s deformation in planar extensional flow: *ε* = (5/2)*μΩ*/*G*, where *ε* = (*a – b*)/(*a + b*) is the now spheroidal object’s strain (long axis *a*, short axes *b* = *c*), *Ω* is the uniform extensional strain rate of the undisturbed flow field, *μ* is the fluid Newtonian viscosity, and *G* is the sphere’s elastic shear modulus. In our L-EV mechanical measurements, *ε* is the average strain of the ellipses fitted to the L-EV contours along its trajectory. We estimated the *Ω* experienced by suspended L-EVs from the microfluidic device dimensions (*h*, *w*) and prescribed flow rates (*Q*) as *Ω* ~ 1.5*U*/(*w*/2) where *U* = *Q*/(*hw*) is the average velocity in the rectangular channels leading the stagnation point region.^[29]^ The viscosity is estimated from rheometer measurements of the suspending fluid (see *Suspending Fluid* for more details).

*Statistical Analysis of Stiffness Distributions:* The number of L-EVs measured in a single experiment is *N*. For the MGG and Gli36 IDH1 wild-type vs. IDH1 mutant comparisons, two independent repeated experiments were conducted. For all other cell lines, one experiment was performed. Statistically significant differences of L-EV stiffness distributions from different cell lines were assessed to a significance level of at least p < 0.05 by comparing the medians using the two-sided Wilcoxon rank sum test (also known as the Mann-Whitney U-test) as implemented in MATLAB. The Wilcoxon rank sum test is a nonparametric test like the one-way ANOVA that is appropriate for distributions that are not normally distributed. One, two, or three asterisks represent significance levels of p < 0.05, p < 0.01, and p < 0.001, respectively. Normality of stiffness distributions was judged from the Lilliefors test, which tests if distributions come from the normal family to 5% significance level.

*Morphology:* Morphology and size distributions of L-EVs were assessed from optical microscopy images acquired during the microfluidic L-EV stiffness measurement experiments. Microscopy images of L-EVs and other aggregates were obtained from the frames of movies captured during extensional flow stretching experiments. Images of quasi-spherical L-EVs that were candidates for the stiffness measurement and non-spherical clumpy aggregates that were not measured for stiffness were captured. The effective diameter *d* of an L-EV was based on the average area 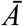 contained in the detected contours along the L-EV’s trajectory: 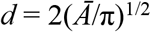.

*L-EV Concentration Estimation:* The concentration of L-EVs suspended in aqueous methyl cellulose in PBS was determined from vertical stacks of 2D microscopy images (z-stack) of the L-EV suspension in the microfluidic device in a stationary state (flow rate 0 μL/hr). Concentration was the ratio of the number of single L-EVs and other aggregates were counted divided by the known z-stack volume. For each sample measured, 4 – 13 z-stacks were averaged. The total number of objects and single near-spherical L-EVs that were suitable for mechanical property measurements were accounted for separately.

*Lipidomics:* Prior to analysis, aliquots of each lipid extract were diluted in isopropanol:methanol (2:1, v:v) containing 20 mM ammonium formate. Full scan MS spectra at 100,000 resolution were collected on a LTQ-Orbitrap Velos mass spectrometer in both positive and negative ionization modes. A Shimadzu Prominance HPLC served as the sample delivery unit. The sample and injection solvent were 2:1 (v: v) isopropanol: methanol containing 20 mM ammonium formate. The spray voltage was 4.5 kV, ion transfer tube temperature was 275 °C, the S-lens value was 50%, and the ion trap fill time was 100 ms. The autosampler was set to 15 °C. Following MS data acquisition, offline mass recalibration was performed with Thermo Xcalibur software according to the vendor’s instructions. MS/MS confirmation and structural analysis of lipid species identified by database searching were performed using higher-energy collisional dissociation (HCD) MS/MS at 60,000 resolution and a normalized collision energy of 25 for positive ion mode, and 60 for negative ion mode.

*Quantification of Lipid Peaks*: Lipids were identified using the Lipid Mass Spectrum Analysis (LIMSA) v.1.0 software linear fit algorithm, in conjunction with an in-house database of hypothetical lipid compounds, for automated peak finding and correction of 13 °C isotope effects. Peak areas of found peaks were quantified by normalization against an internal standard of a similar lipid class. The top ~300 most abundant peaks in both positive and negative ionization mode were then selected for MS/MS inclusion lists and imported into Xcalibur software for structural analysis on the pooled QC sample as described above.

*Analysis of Lipidomics:* The absolute values of lipids were divided with the total lipids amount to present the portion of them. To show the altered expression of lipids, heatmap and graphs were attained by Prism 9 software. For principal component analysis (PCA), all the lipid portion data was processed using R program (version 4.0.3).

## Supporting Information

Supporting Information is available from the Wiley Online Library and from the authors upon request.

## 6. Acknowledgements

This work was supported in part by National Institute of Health grants (R21CA217662, R01GM138778 to H.I.). M.H.J. is supported by Basic Science Research Program through the National Research Foundation of Korea (NRF) funded by the Ministry of Education (NRF-2021R1A6A3A14039686).

