## Supplementary Information for "Non-contact microfluidic analysis of the stiffness of single large extracellular vesicles from IDH1-mutated glioblastoma cells"

### Sources of Uncertainty in L-EV Effective Stiffness Measurements

#### *Microfluidic Flow Field Deforming the L-EVs*

The flow conditions experienced by the deforming L-EVs were estimated in a consistent manner across all experiments. The absolute value of the reported stiffnesses was affected by these estimates, but all by the same amount. The first approximation was the estimation of the strain rate of the extensional flow in the stagnation point region of the microfluidic device according to our previous study.[29] This extensional strain rate estimate depended on device width and height and flow rate. The same device dimensions and range of flow rates were used for all experiments. Thus, the systematic error introduced by the strain rate estimate was the same for all experiments. The second approximation was that L-EVs were deformed in perfectly planar extensional flow. However, L-EVs were imaged at 35 – 40 µm above the device's glass bottom surface, that is less than the 75-µm mid-channel height. This was the furthest distance possible with the 100x objective used that maintained good image quality. This close to the wall, the parabolic velocity profile was still changing with vertical distance from the glass bottom, meaning that the flow was not perfectly planar as assumed by the mechanical model. However, the variation in velocity across the diameter of the very small L-EVs (~1 – 3.5 µm diameter) is <10%.[57] Considering that all L-EVs were consistently measured in a narrow range 35 – 40 µm above the glass bottom, the introduced error was similar across all stiffness measurements. Therefore, we do not attribute the fluid strain rate and planar flow approximations to be major causes of the discrepancy between Figure 4A and Figure 4B because the device dimensions, flow rates, and imaging plane were very similar. The precision of our microfluidic L-EV stiffness measurements could be improved by using a more sophisticated simulation to determine the fluid strain rates experienced by L-EVs. A finite element software could simulate the strain rates experienced by L-EVs that are 35-40 µm above the bottom surface. This change could help to increase the measurement accuracy and precision.

*Suspending fluid: Aqueous methyl cellulose in phosphate buffered saline*

The 0.6% w/v and 0.7% w/v aqueous methyl cellulose in PBS solutions used in the measurements were shear thinning (**Figure S4**). This meant that viscosity decreased with increasing shear rate. In the microfluidic device, the deforming fluid shear stress was in the range 2.8 – 8.3 $s^{-1}$. However, the rheometer's viscosity measurement readings in this low shear rate range were unstable and noisy. Due to this instrument limitation, the estimate of viscosity used in the analytical mechanical model was the average viscosity across a range of shear rates higher shear rates, typically over 5 – 50 $s^{-1}$. Given that aqueous methyl cellulose is shear thinning, the viscosity was an underestimate of what the L-EVs actually experienced. Furthermore, the viscosity experienced by L-EVs was not constant over its surface because the flow field was not perfectly planar. The faster velocities and higher shear rates on the top sides of the L-EVs (closer to the mid-channel height) resulted in lower viscosities in this area compared to the bottom sides (closer to the glass bottom). The compounding error effects of the shear thinning fluid and varying shear rates likely made the viscosity the quantity with the highest uncertainty in the analytical mechanical model. Therefore, when the viscosity was changed it was not surprising to have the absolute L-EV effective stiffnesses not match between the Gli36 and MGG WT vs. IDH1 mutation independent experiments shown in Figure 4A and Figure 4B.

**Supplementary Tables**

| Cell Line | *N* | *G* Distribution Mean ± SEM [Pa] | *G* Distribution Median [Pa] | Fluid Viscosity [mPa·s] | p-value |
|---|---|---|---|---|---|
| T98G | 56 | 4.82 ± 0.33 | 4.25 | 30 | 0.4519 |
| Gli36 WT | 118 | 5.80 ± 0.36 | 4.50 | | |
| T98G | 56 | 4.82 ± 0.33 | 4.25 | 30 | 0.004464 |
| MGG WT | 90 | 6.17 ± 0.33 | 5.84 | | |
| T98G | 56 | 4.82 ± 0.33 | 4.25 | 30 | 0.004358 |
| U87 | 78 | 6.32 ± 0.37 | 5.85 | | |
| Gli36 WT | 118 | 5.80 ± 0.36 | 4.50 | 30 | 0.09665 |
| MGG WT | 90 | 6.17 ± 0.33 | 5.84 | | |
| Gli36 WT | 118 | 5.80 ± 0.36 | 4.50 | 30 | 0.08454 |
| U87 | 78 | 6.32 ± 0.37 | 5.85 | | |
| MGG WT | 90 | 6.17 ± 0.33 | 5.84 | 30 | 0.9126 |
| U87 | 78 | 6.32 ± 0.37 | 5.85 | | |
| Gli36 WT | 118 | 5.80 ± 0.36 | 4.50 | 30 | 0.0004636 |
| Gli36 IDH1 mut | 142 | 7.53 ± 0.38 | 6.41 | | |
| MGG WT | 90 | 6.17 ± 0.33 | 5.84 | 30 | 0.01303 |
| MGG IDH1 mut | 245 | 7.64 ± 0.28 | 7.32 | | |
| Gli36 WT | 203 | 14.72 ± 0.60 | 12.42 | 65 | 0.02048 |
| Gli36 IDH1 mut | 172 | 16.88 ± 0.73 | 15.14 | | |
| MGG WT | 198 | 14.25 ± 0.60 | 12.52 | 68 | 0.006032 |
| MGG IDH1 mut | 190 | 17.23 ± 0.75 | 14.65 | | |

**Table S1.** Summary of hypothesis testing statistics for comparisons between L-EV stiffness distributions in Figures 3 and 4. The two-sided Wilcoxon rank sum test was used. The null hypothesis was that the two data sets come from distributions with equal medians. Distribution differences were considered statistically significant for $p < 0.05$. In these cases, the null hypothesis was rejected, meaning that these two data sets came from distributions with different medians. *N* was the number of L-EV stiffness measurements in each distribution. SEM was standard error of the mean. The viscosity was the viscosity of the fluid that L-EVs were suspended in for the microfluidic stiffness measurements.

### Supplementary Figures

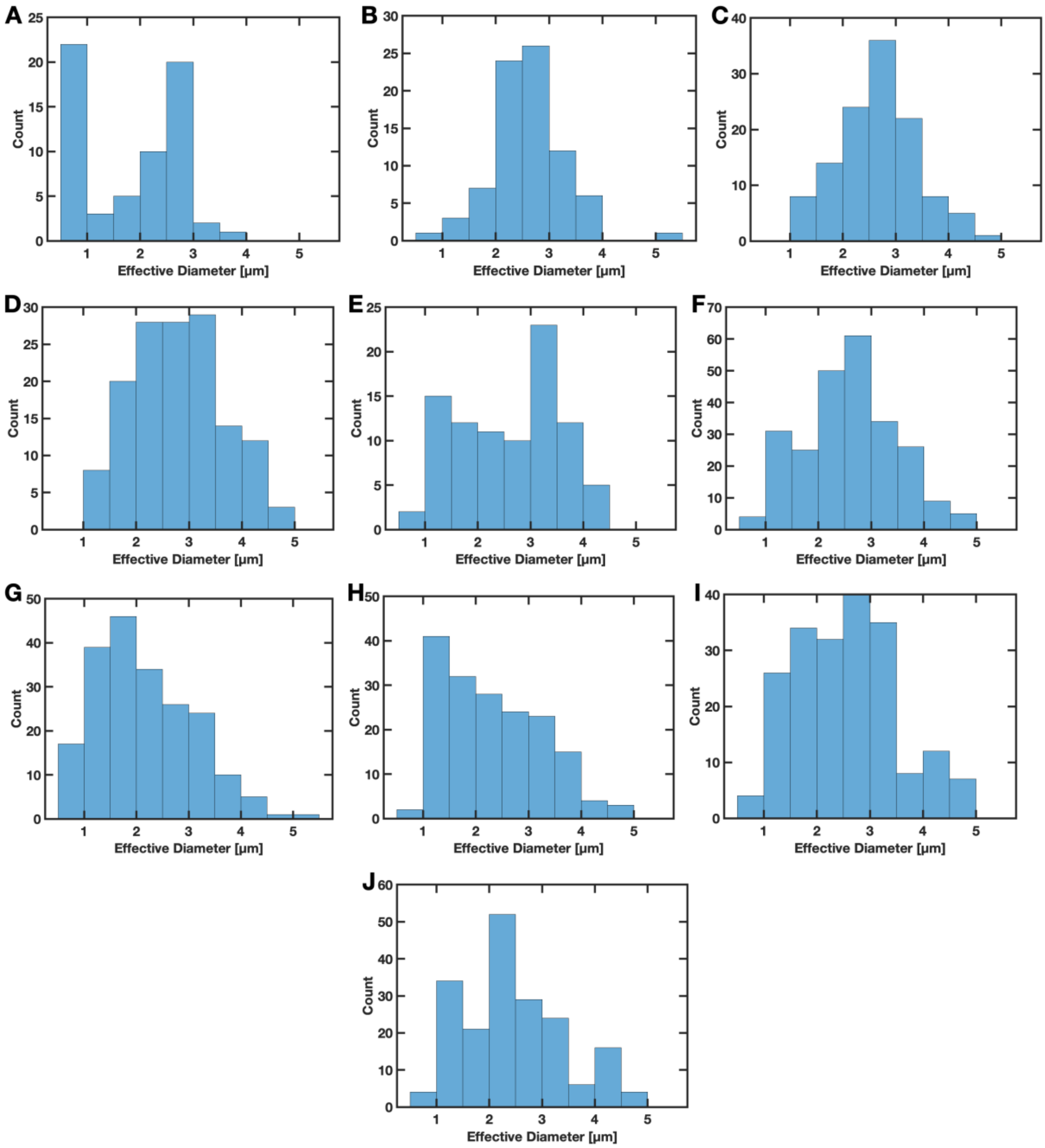


**Figure S1.** Size distributions of large extracellular vesicles measured for each glioblastoma cell line. **(A)** T98G, **(B)** U87 **(C)** Gli36 wild-type, fluid viscosity $\mu$ ~ 30 mPa·s, **(D)** Gli36 IDH1 mutant, $\mu$ ~ 30 mPa·s, **(E)** MGG wild-type, $\mu$ ~ 30 mPa·s, **(F)** MGG IDH1 mutant $\mu$ ~ 30 mPa·s, **(G)** Gli36 wild-type, $\mu$ ~ 65 mPa·s, **(H)** Gli36 IDH1 mutant, $\mu$ ~ 65 mPa·s, **(I)** MGG wild-type, $\mu$ ~ 68 mPa·s, **(J)** MGG IDH1 mutant, $\mu$ ~ 68 mPa·s.

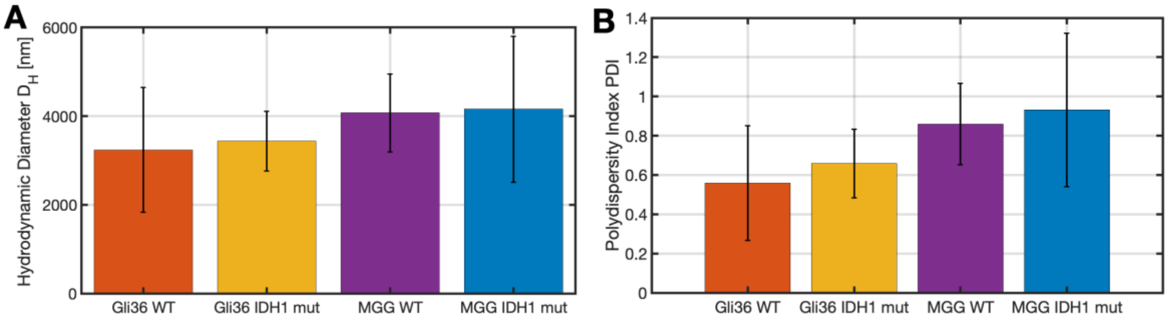


**Figure S2.** Large extracellular vesicle size characterization by dynamic light scattering (DLS). **(A)** Hydrodynamic diameter $D_H$. **(B)** Polydispersity index.

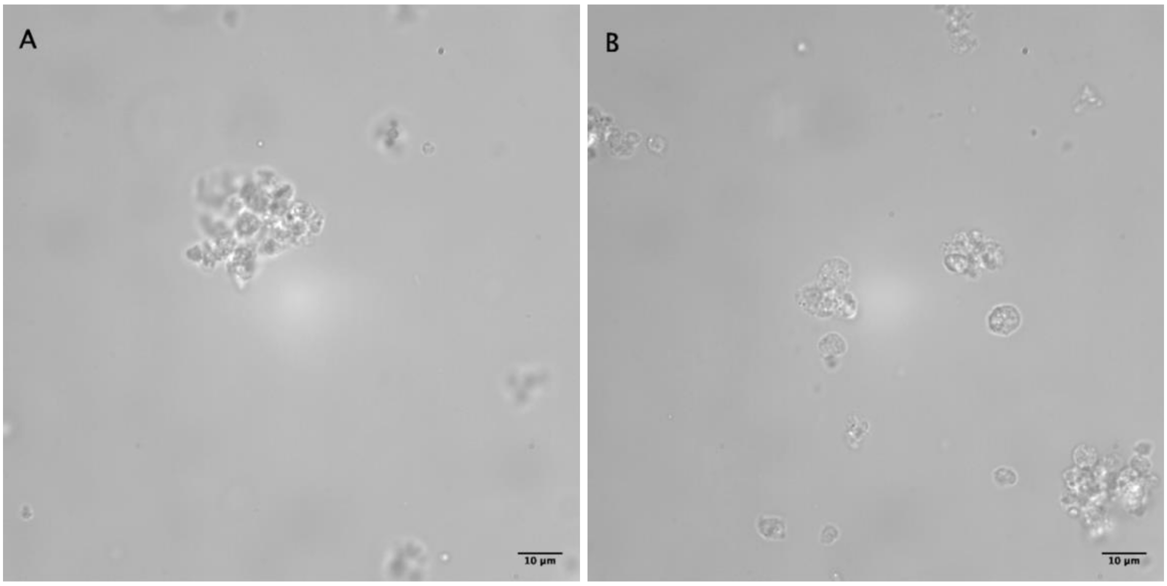


**Figure S3.** Representative images of large extracellular vesicle aliquot contents in a fluid chamber with no fluid flow. A defrosted 20-µL aliquot derived from T98G cells was suspended in 0.6% w/v aqueous methyl cellulose in phosphate buffered saline. Suspensions were imaged in a CoverWell perfusion chamber gasket (Invitrogen, C18139, 9 mm diameter, 0.5 mm deep). Large aggregates greater than ~10 µm in size were common while isolated <5 µm spherical L-EVs were difficult to find.

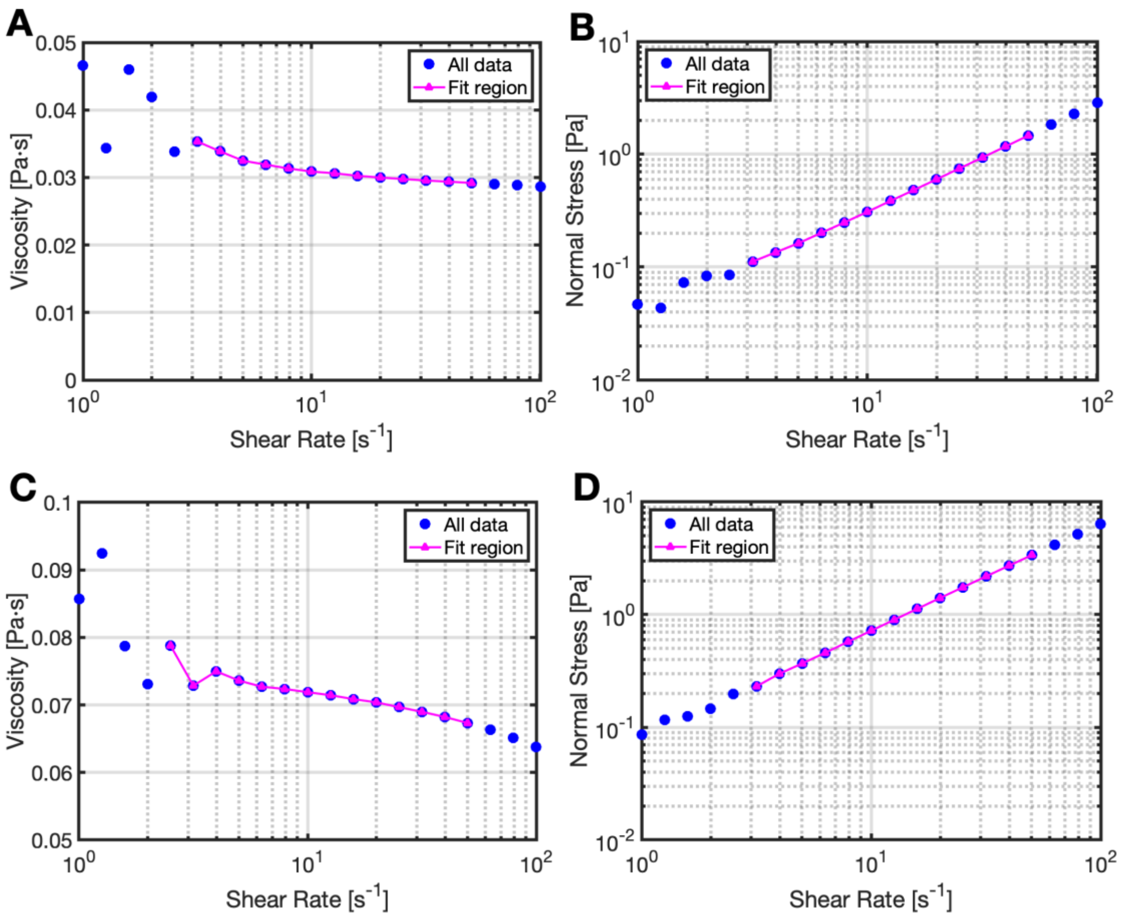


**Figure S4.** Representative flow sweep measurement data on a rheometer for aqueous methyl cellulose in phosphate buffered saline (mcPBS). The viscosity was estimated to be the average viscosity in a low shear rate regime (fit region) after initial transients had died out and the normal force was linear. The viscosity used in the analytical technical model was the average of three independent viscosity measurements. 0.6% w/v mcPBS **(A)** viscosity and **(B)** normal stress. 0.7% w/v mcPBS **(C)** viscosity and **(D)** normal stress. Lines between data points in the fit region are guides for the eye to show data trends.

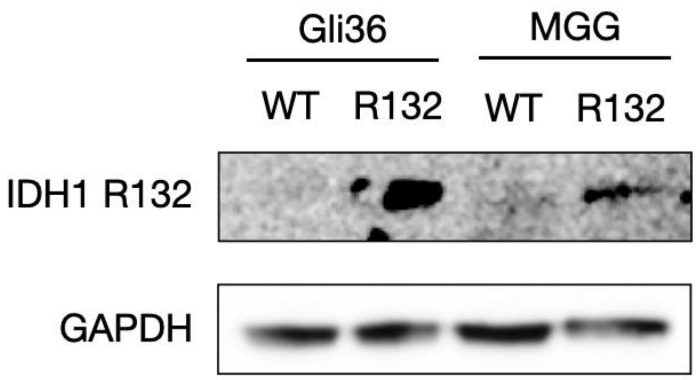


**Figure S5.** The expression of the mutated IDH1 R132 in wild-type of Gli36 and MGG cells, and IDH1-mutated Gli36 and MGG.

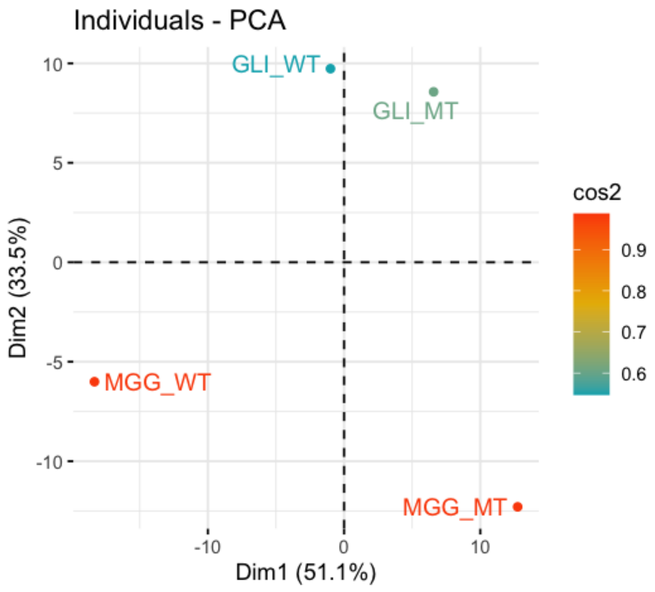


**Figure S6.** Principal component analysis (PCA) of lipid composition of L-EVs from Gli36 and MGG cell lines in wild-type and IDH1 mutation.

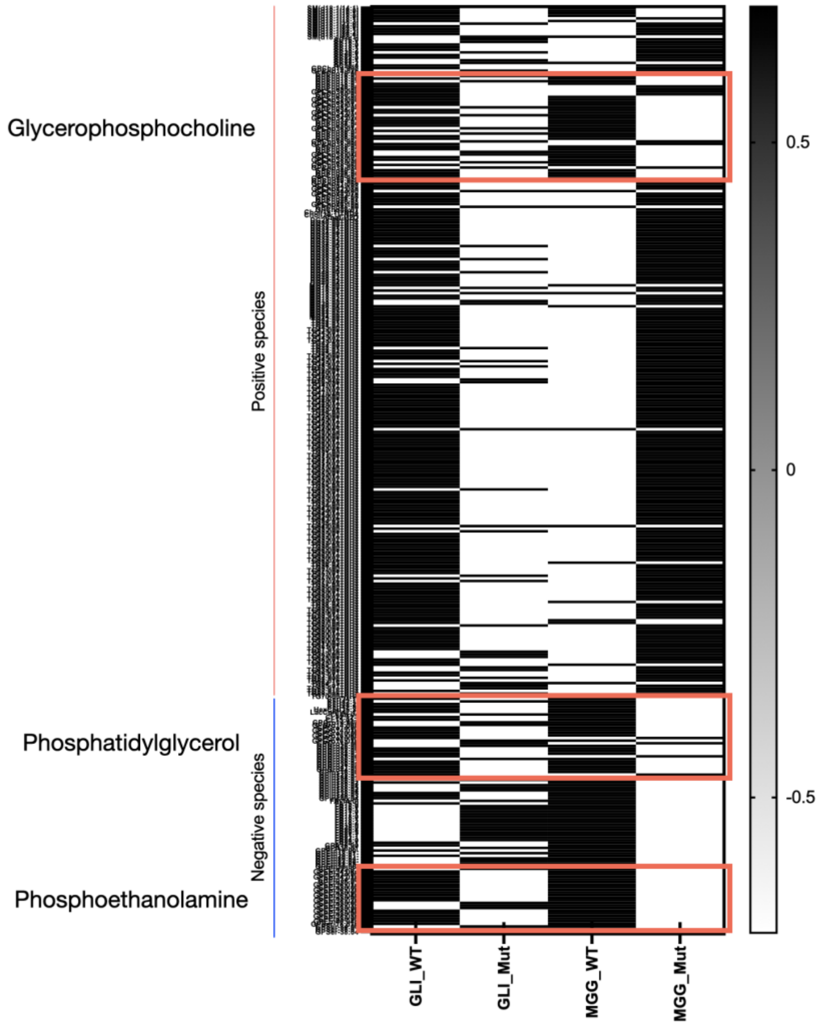


**Figure S7.** The heatmap of lipid profiling in apoptotic bodies from IDH1 wild-type and IDH1-mutated cells.
